# Effect of Adhesion and Substrate Elasticity on Neutrophil Extracellular Trap Formation

**DOI:** 10.1101/508366

**Authors:** Luise Erpenbeck, Antonia Luise Gruhn, Galina Kudryasheva, Gökhan Günay, Daniel Meyer, Elsa Neubert, Julia Grandke, Michael P. Schön, Florian Rehfeldt, Sebastian Kruss

## Abstract

Neutrophils are the most abundant type of white blood cells. Upon stimulation, they are able to decondense and release their chromatin as neutrophil extracellular traps (NETs). This process (NETosis) is part of immune defense mechanisms but also plays an important role in many chronic and inflammatory diseases such as atherosclerosis, rheumatoid arthritis, diabetes and cancer. For this reason, much effort has been invested into understanding biochemical signaling pathways in NETosis. However, the impact of the mechanical micro-environment and adhesion on NETosis is not well understood.

Here, we studied how adhesion and especially substrate elasticity affect NETosis. We employed polyacrylamide (PAA) gels with distinctly defined elasticities (Young’s modulus *E)* within the physiologically relevant range from 1 kPa to 128 kPa and coated the gels with integrin ligands (collagen I, fibrinogen). Neutrophils were cultured on these substrates and stimulated with potent inducers of NETosis: phorbol 12-myristate 13-acetate (PMA) and lipopolysaccharide (LPS). Interestingly, PMA-induced NETosis was neither affected by substrate elasticity nor by different integrin ligands. In contrast, for LPS stimulation, NETosis rates increased with increasing substrate elasticity (*E* > 20 kPa). LPS-induced NETosis increased with increasing cell contact area, while PMA-induced NETosis did not require adhesion at all. Furthermore, inhibition of phosphatidylinositide 3 kinase (PI3K), which is involved in adhesion signaling, completely abolished LPS-induced NETosis but only slightly decreased PMA-induced NETosis.

In summary, we show that LPS-induced NETosis depends on adhesion and substrate elasticity while PMA-induced NETosis is completely independent of adhesion.

## Introduction

Neutrophilic granulocytes are the most abundant type of circulating white blood cells. In a process termed NETosis, they release neutrophil extracellular traps (NETs), web-like structures composed of decondensed chromatin and decorated with antimicrobial proteins^1,2^. During NETosis, the nuclear chromatin swells until both the nuclear envelope and the outer cell membrane rupture^3^. NETosis is considered an important immune defense mechanism as neutrophils can bind and kill bacteria and other pathogens via NETs. Apart from physiological stimuli such as pathogens, chemokines (e.g. CXCL8), activated platelets or urea crystals there are several additional NET-inducers like phorbol 12-myristate 13-acetate (PMA) and lipopolysaccharides (LPS), which induce NETosis *in vitro*^4^. Even though NETosis was initially described as part of the innate immune defense system, we know today that dysregulated NETosis is also involved in a variety of chronic inflammatory and autoimmune diseases such as atherosclerosis, systemic lupus erythematosus, preeclampsia, as well as malignant diseases^5–8^. Therefore, the question which environmental factors play a role in this process and may influence the course of diseases is highly important.

Mechanical properties of tissues are environmental signals that are able to modulate the functionality of surrounding cells. This has been demonstrated by a substantial amount of studies investigating the effect of physical factors on cellular functions^9,10–12^. It has previously been shown that phenotype and functionality of immune cells such as macrophages and dendritic cells are affected by substrate elasticity/stiffness^13–15^. It has also been reported that substrate elasticity affects neutrophil adhesion, migration and chemotaxis^16–18^. Transmigration of neutrophils through endothelium was also proven to be affected by sub-endothelial cell matrix stiffness^19^. Tissue stiffness increases in multiple pathological processes including, most prominently, atherosclerotic plaques^20^ but also fibrosis^21^ and cancer^22^. In general, cell adhesion is mediated through surface receptors interacting with specific ligands presented on surfaces^23–25^. Integrin ligands have been previously shown to play an important role in leukocyte adhesion and migration^26–28^. Additionally, the ligand density on the surface affects adhesion and migration of neutrophils^28,29^. For example, neutrophils adhere *via* the integrin Mac-1 to the platelet receptor GPIbα and show the fastest adhesion maturation when ligands where distributed in a medium distance of approximately 100 nm^28^. Even though the role of integrins on neutrophil adhesion has been studied, there are contradictory studies concerning their role in NETosis^30,31^. Exemplarily, it has been shown that blocking of the integrin LFA-1 prevented NETosis^32^. However, the overall impact of substrate elasticity and adhesion-related processes on NETosis remain poorly understood.

In this paper, we explore the effect of substrate stiffness/elasticity (Young’s modulus *E*) and general adhesion on NETosis in human neutrophils. We use collagen I- and fibrinogen-coated polyacrylamide (PAA) gels with stiffnesses within the physiologically relevant range (*E* = 1 kPa to 128 kPa) and study the impact of elasticity and adhesion on NETosis induced by two different stimuli (LPS, PMA).

## Material & Methods

### Polyacrylamide (PAA) gel preparation

PAA gels were prepared on round glass cover slides as previously reported^9,10^. In brief, the cover glasses were plasma cleaned, coated with 3-aminopropyltriethoxysilane (Sigma A3648) and afterwards incubated with glutaraldehyde solution (0.05%, Sigma G7651). Appropriate mixtures of acrylamide (Bio-Rad #161-0140) and bis-acrylamide (Bio-Rad #161-0140) diluted in Dulbecco’s phosphate-buffered saline (PBS, Sigma-Aldrich) were freshly prepared, stored at +4 °C and used within 2 months (see table T1 in SI for details). Polymerization was initiated by addition of 1/1000 N,N,N,N-tetramethylethylenediamine (TEMED) and 1/100 ammonium persulfate (APS) solution. 35 µL of this solution was used per cover slip. A square hydrophobic cover glass was placed on top in order to equally distribute the solution on the bottom glass. Gels were polymerized for 60 minutes in a saturated water atmosphere to avoid evaporation and were approximately 70 µm thick. Young’s modulus *E* of PAA gels was quantitatively controlled by measurements on a bulk rheometer (MCR-501, Anton Paar, Austria). To prevent toxicity to cells, non-polymerized residues were thoroughly washed away using PBS. Substrates were treated with the photo-activatable cross-linker Sulfo-SANPAH (Thermo Scientific 22589; 0.4 mM in 50 mM HEPES buffer at pH 8), exposed to UV light (λ = 365 nm) for 10 minutes and then either coated with rat tail collagen I (0.02 mg/mL for data presented in all main figures and 0.2 mg/mL for supplementary figure S3) (Corning #354236) or fibrinogen (0.02 mg/mL, from human plasma, Sigma # F-3879) overnight at 4 °C unless otherwise stated.

### Human neutrophil isolation

All experiments with human neutrophils were approved by the Ethics Committee of the University Medical Center (UMG) Göttingen (protocol number: 29/1/17). Donors gave informed voluntary consent to the study. Neutrophils were isolated according to published standard protocols^3,33^ from healthy donors’ venous blood supplemented with EDTA.

In short, fresh blood from donors was collected with S-Monovettes KE 7.5 ml (Sarstedt). The blood was gently layered on top of Histopaque 1119 (Sigma-Aldrich) in a 1:1 ratio and centrifuged at 1100 x g for 21 minutes. Cells were separated by their density resulting in different cell layers being formed. The third (transparent) and forth (pink) layer containing white blood cells were taken and mixed into Hank’s Balanced Salt Solution (HBSS) (without Ca^2+^/Mg^2+^, Thermo Fisher Scientific), followed by centrifugation at 400 x g for 10 minutes. The supernatant was discarded and the pellet was resuspended in HBSS and then layered on top of a phosphate buffered Percoll (GE Healthcare) gradient at concentrations of 85%, 80%, 75%, 70% and 65% and centrifuged at 1100 g for 22 minutes. We collected the layers containing neutrophils and replenished the volume with HBSS and centrifuged at 400 g for 10 minutes. The resulting pellet was resuspended in 1 mL of HBSS and cells were counted using a Neubauer chamber (OptikLabor). The desired cell number was resuspended in Roswell Park Memorial Institute (RPMI) 1640 medium (Lonza) containing 0.5% fetal calf serum (FCS; Biochrom GmbH). Neutrophil purity was determined by a cytospin assay (Shanson, Cytospin 2 Centrifuge) followed by Diff Quick staining (Medion Diagnostics). In all experiments neutrophil purity was >95%.

### Quantification of Neutrophil extracellular trap formation

Freshly isolated neutrophils were seeded (0.5 × 10^6^ cells/well) on PAA gels and collagen I- or fibrinogen-coated glass and incubated for 30 minutes at 37 °C, 5% CO_2_ in order to allow the cells to adhere. NETosis was either stimulated with 5 nM Phorbol-12-myristate-13-acetate (PMA, Sigma-Aldrich) or with 75 μg/mL lipopolysaccharide (LPS) from *Pseudomonas aeruginosa* (serotype 10.22, strain: ATCC 27316, Sigma-Aldrich) and incubated for 3 hours (37 °C, 5% CO_2_). NETosis was stopped with 2% paraformaldehyde (PFA, Roth) fixation overnight. The next day, the fixed neutrophil chromatin was stained using 1 µg/ml Hoechst 33342,trihydrochloride,trihydrate (Hoechst) (life technologies). Cells were imaged with an Axiovert 200 microscope (16x magnification, Zeiss) being set on the blue channel (Filter set49 DAPI shift free, 488049-9901-000, Zeiss) connected to a CoolSNAP ES camera (Photometrics). 6 images of different locations were taken per well. Phase contrast images for cell contact area measurements were taken for each setting and the cell contact areas were calculated by using ImageJ 4.46r for at least 50 cells per condition. NETosis was quantified in a standardized blinded fashion as percentage of condensed and decondensed nuclei.

### PI3K Inhibition

Human neutrophils were treated with the PI3K inhibitor BAY 80-6946 (copanlisib) for 20 minutes on ice and then seeded on PAA gels (4 kPa, 128 kPa) and glass followed by a 30 minute incubation (37 °C, 5% CO_2_). Afterwards cells were stimulated, fixed and imaged as described above.

### Neutrophil extracellular trap formation on passivated glass surfaces

Human neutrophils (10.000 cells/well) were seeded on glass surfaces coated with 0.5 mg/mL Poly-L-lysine (PLL, Sigma) or Poly-L-lysine-grafted-polyethylenglycol (PLL-g-PEG (SuSoS Surface Technology) and incubated for 30 minutes as described before^28^. Uncoated glass surfaces were used as controls. Then cells were stimulated, fixed and stained as described above. For Reflection Interference Contrast Microscopy (RICM) a special objective was used (63x magnified EC Plan-Neofluar Ph3 objective/420481-9911-000, 1.6x Optovar, Zeiss) and also a RICM filter set (reflector module Pol ACR P&Cfor HBO 100/ 424924-9901-000 and emission filter 416 LP, AHF-Nr.: F76-416/ 000000-1370-927, Zeiss).

## Results

### Substrate elasticity affects NETosis in a stimulant-dependent manner

To investigate the effect of substrate elasticity/stiffness on NETosis, freshly isolated human neutrophils were seeded on PAA gels coated with either collagen I or fibrinogen which are both known to interact with integrins on neutrophils^29,34^. The PAA substrate elasticity was varied within the physiological range (1 kPa, 2 kPa, 4 kPa, 8 kPa, 16 kPa, 20 kPa, 30 kPa, 64 kPa, 128 kPa). The prepared gels cover a wide range of physiological elasticities for example that of brain tissue (< 1 kPa), muscle (14-16 kPa), osteoids, pre-calcified bone (30 kPa) or cartilage (>100 kPa).^9^ The coating procedure leads to a uniform high density distribution of the proteins on the surface^10,35^. To understand how substrate elasticity and the presence of certain integrin ligands affect NETosis, neutrophils were seeded and incubated for 30 minutes and then stimulated with either PMA or LPS for 3 hours **(figure 1)**. PMA is a well-known activator of protein kinase C (PKC) and frequently used to induce NETosis *in vitro*^36^. LPS on the other hand, induces NETosis in a receptor-mediated fashion^37^. PMA was applied at a very low concentrations (5 nM) to avoid the strong stimulation at typical concentrations (100 nM^3^) that could blur the impact of substrate elasticity.

**Figure 1.**
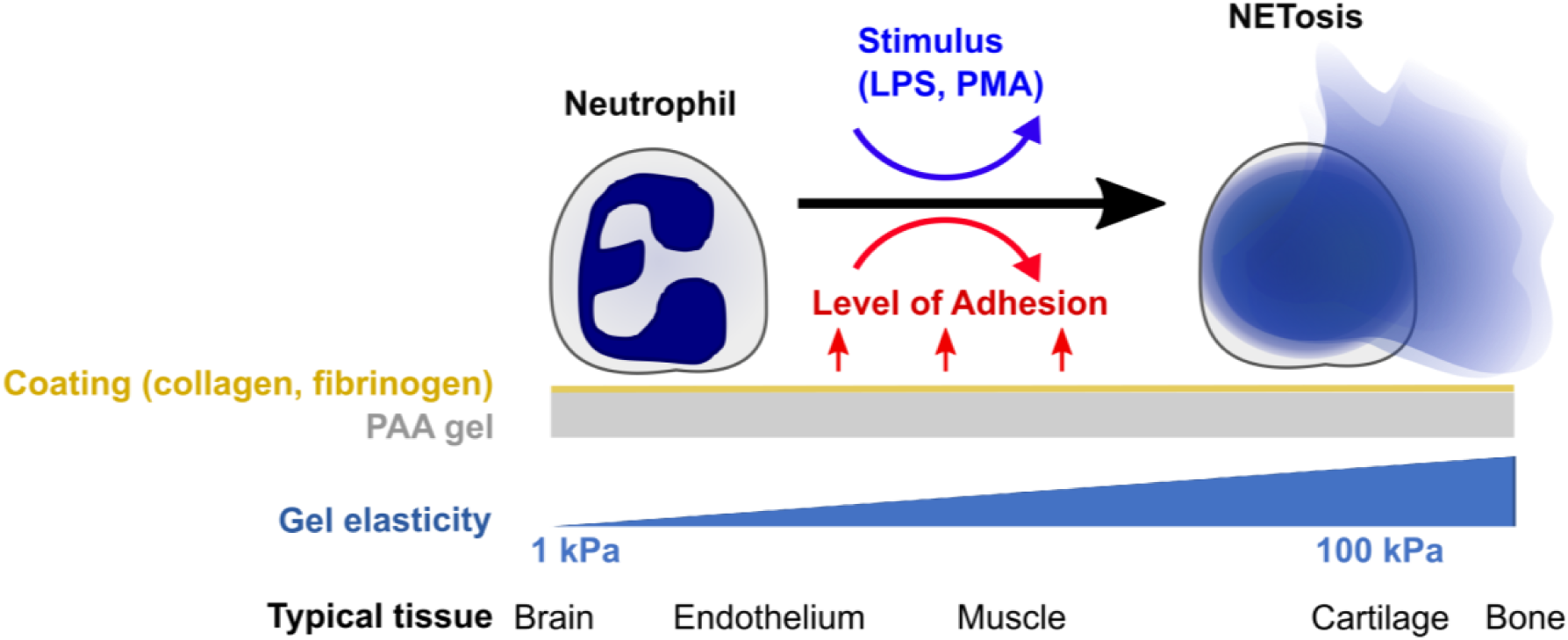
Quantifying the impact of substrate elasticity, adhesion and stimulation on NETosis. Human neutrophils are cultured on polyacrylamide (PAA) gels of different elasticity/stiffness and coating to control and vary adhesion. Cells are then stimulated with PMA or LPS to assess the impact of the different environmental factors on NETosis.

NETosis was imaged and quantified on both surface coatings (collagen I and fibrinogen) for both ways of stimulation (PMA and LPS) **(figure 2)**. Representative images for 2 kPa, 16 kPa, 128 kPa PAA gels and a glass control are shown in **figure 2**.

**Figure 2.**
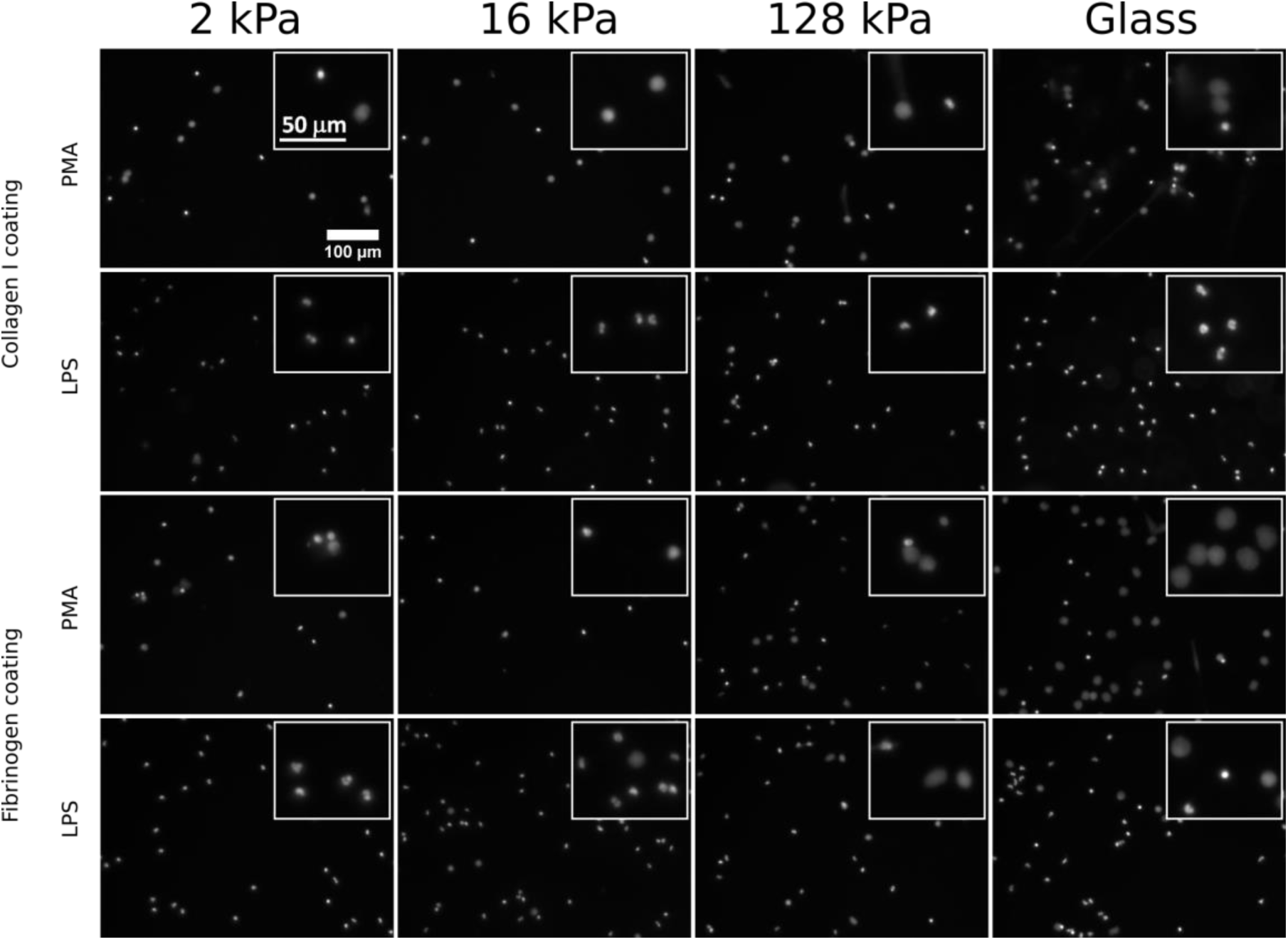
Substrate elasticity affects NETosis in a stimulant-dependent manner. Human neutrophils were seeded on PAA gels coated with collagen I or fibrinogen. They were stimulated with either PMA (5 nM) or LPS (75 μg/mL) as indicated, and incubated for 3h. The fluorescence images show the nuclei (Hoechst DNA/chromatin stain) of fixed cells. Insets show magnified cells/nuclei. The images show that NETosis depends on all three parameters (substrate elasticity, stimulant, surface coating). See quantification in figure 3.

Images of all other conditions are shown in the **supplementary figures S1 and S2**. PMA-stimulated NETosis was independent of stiffness. On collagen I-coated substrates, the NETosis rate was the same for stiffness values **(figure 3a)**. Similarly, on fibrinogen-coated substrates, PMA-induced NETosis **(figure 3c)** did vary with stiffness. However, on glass surfaces NETosis was significantly higher compared to that on substrates with defined stiffnesses of 8 kPa, 16 kPa, 20 kPa or 128 kPa, respectively. This observation might be explained by different effective concentrations of PMA due to diffusion into the gel or adsorption onto glass. We concluded that PMA-induced NETosis is not affected by substrate stiffness.

**Figure 3.**
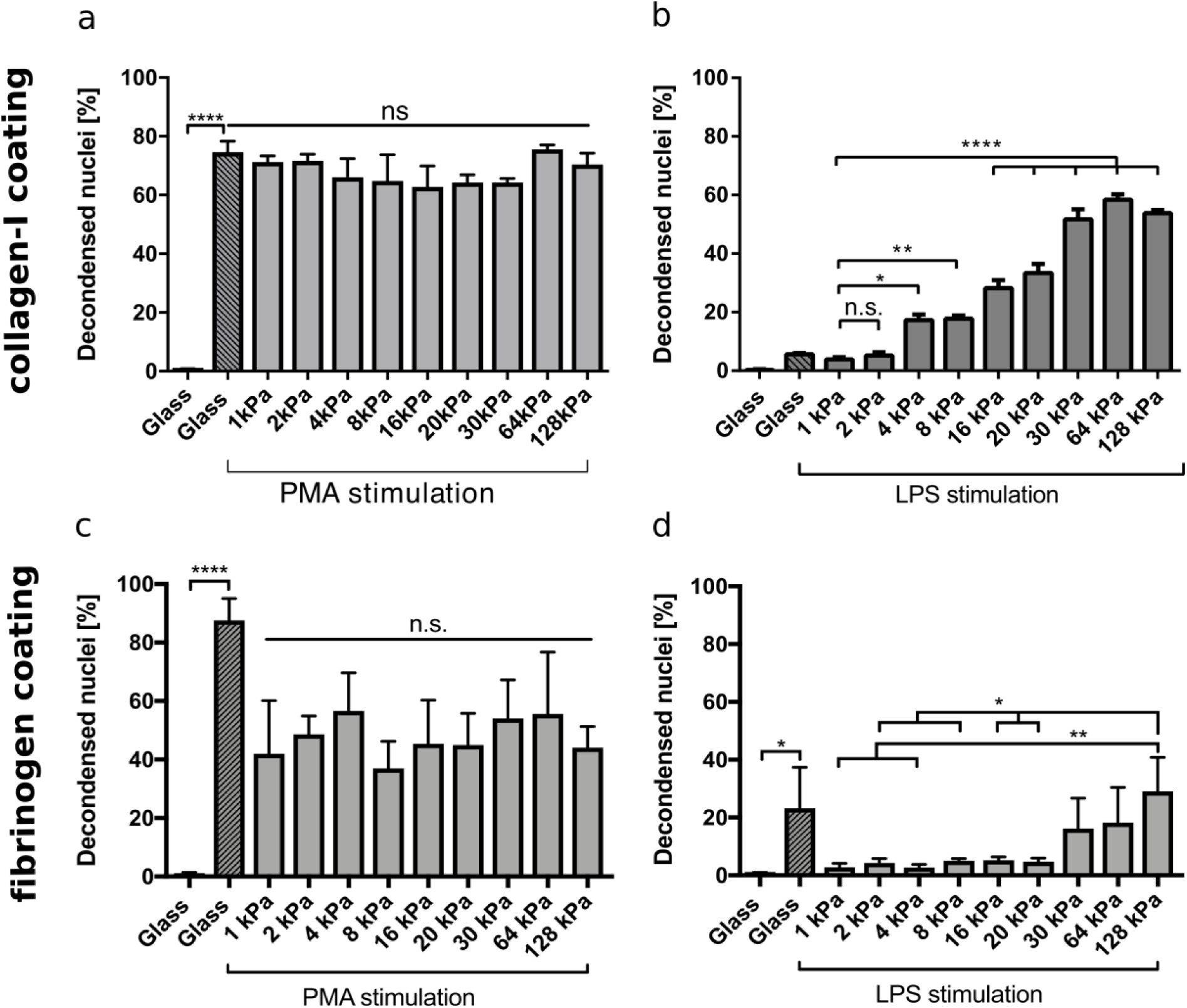
NETosis depends on substrate elasticity, stimulant and surface coating. Human neutrophils were seeded on PAA gels and glass surfaces coated with either collagen I (a, b) or fibrinogen (c,d) and stimulated with either PMA (5 nM)(a, c) or LPS (75 μg/mL)(b, d) and incubated for 3h. PMA-induced NETosis does not depend on substrate elasticity. LPS-induced NETosis increases significantly for both coatings above a stiffness threshold of >20 kPa. n > 500 cells for each condition. N = 3 donors. Statistics: one-way ANOVA (Bonferroni’s multiple comparisons test; * p < 0.05; ** p < 0.01; **** p < 0.0001; ns: not significant). Mean ± SEM.

In contrast, NETosis was significantly affected by substrate elasticity under LPS stimulation on both collagen and fibrinogen-coated surfaces **(figure 3b,d)**. On stiffer substrates (E > 20 kPa) NETosis was significantly increased.

### Correlation between neutrophil adhesion and NETosis

To further investigate the connection between NETosis, adhesion and substrate elasticity we assessed neutrophil adhesion on the different substrates. Although it is known that upon stimulation with PMA or LPS neutrophils initially adhere strongly to the provided surfaces before starting NETosis^3^, it remains controversial whether this initial adhesion is a necessary prerequisite and to which extent NETosis and adhesion are correlated with one another. During NETosis, the cells round up and the cytoskeleton is degraded. There have been contradictory studies on the role of integrin receptor engagement during NET formation^30,31^. The substrate stiffness most likely influences adhesion. Indeed, neutrophils have been reported to better adhere to stiff surfaces compared to soft surfaces^16,17^. Therefore, we next analyzed the spreading of human neutrophils on collagen I- and fibrinogen-coated substrates and investigated the correlation between adhesion and NETosis. As a surrogate parameter for adhesion we determined the cell contact area (by phase contrast microscopy). On both fibrinogen and collagen I-coated PAA gels the cell contact area/spreading area of neutrophils increased with increasing stiffnesses (**figure 4a,b**), corresponding to the measured LPS-induced NETosis rates **(figure 3 b,d)**. Next, a cross-correlation analysis for neutrophil spreading area versus NETosis was carried out for LPS stimulation. LPS-induced NETosis correlated well with spreading area for both coatings **(figure 4 c,d)**. Therefore, adhesion signaling appears to be highly relevant for LPS-induced NETosis. Differences in adhesion due to different stiffnesses directly translate into different NETosis rates.

**Figure 4.**
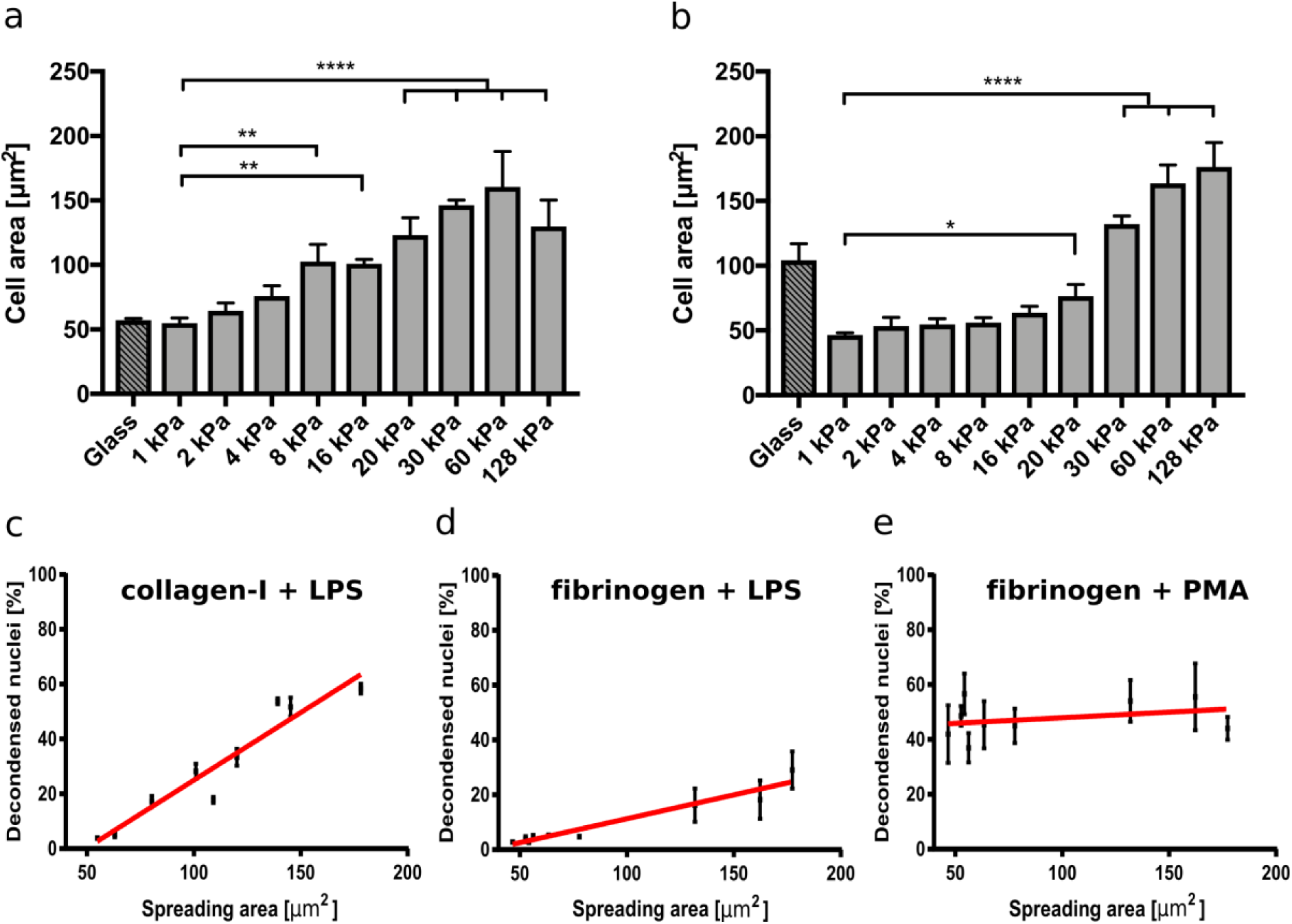
NETosis correlates with cell spreading area. (a) Cell spreading area increases with increasing stiffness on collagen I-coated PAA gels. (b) Cell spreading area increases with increasing stiffness on fibrinogen-coated PAA gels. LPS-stimulated NETosis correlates with spreading area on PAA gels of different stiffness coated with collagen (c) and fibrinogen (d). (e) PMA stimulation does not depend on spreading area. The red lines indicate linear fits. R^2^=0.91 (a). R^2^=0.67 (b). R^2^=0.02 (c). n > 500 cells for each condition. N = 3 donors. Mean ± SEM. Statistics: one-way ANOVA (Bonferroni’s multiple comparisons test. * p < 0.05; ** p < 0.01; **** p < 0.0001; ns: not significant).

In contrast, as expected from the results in figure 3, PMA-induced NETosis did not correlate with the spreading area (**figure 4e).** Interestingly, when the gels were coated with a tenfold higher (0.2 mg/ml) collagen I concentration, cells did not perform NETosis at all (**supplementary figure S3**). We hypothesized that there is an optimum density for NET formation on collagen I, in accordance with previously published results that show an optimum ligand-receptor ratio for different biological functions including cell adhesion or spreading^28,38–40^. Similarly, the cells did not adhere well and did not spread above the area expected from a fully settled neutrophil (**supplementary figure S3**). These observations additionally corroborate our hypothesis that conditions that affect the adhesive phenotype change NETosis rates, which suggests that adhesion and NETosis are interconnected.

### PI3K inhibition abrogates stiffness-dependent variations in NETosis

PI3K activity is important for neutrophil mechanosensing and enables these cells to distinguish between substrates of different stiffnesses^16^. To corroborate that the observed effects on LPS-induced NETosis could be attributed to variations in substrate stiffness, we treated neutrophils with a potent PI3K inhibitor, BAY 80-6946 (copanlisib), and analyzed adhesion and NETosis. Fibrinogen-coated PAA gels of 4 kPa and 128 kPa, respectively, were chosen as representative conditions for adhesion and NETosis quantification as they had revealed significant differences in the experiments presented above. Adhesion was impaired when neutrophils on stiff surfaces (128 kPa and glass) were pretreated with the inhibitor **(figure 5a)**. PI3K is a known signaling intermediate for PMA-induced NETosis, which is why moderate effects on PMA-induced NETosis were to be expected^37^. To investigate the connection between adhesion and NETosis, we treated cells with PMA or LPS under PI3K inhibition. This treatment partially impaired PMA-induced **(figure 5b)**, and completely abrogated LPS-induced NETosis **(figure 5c)**.

**Figure 5.**
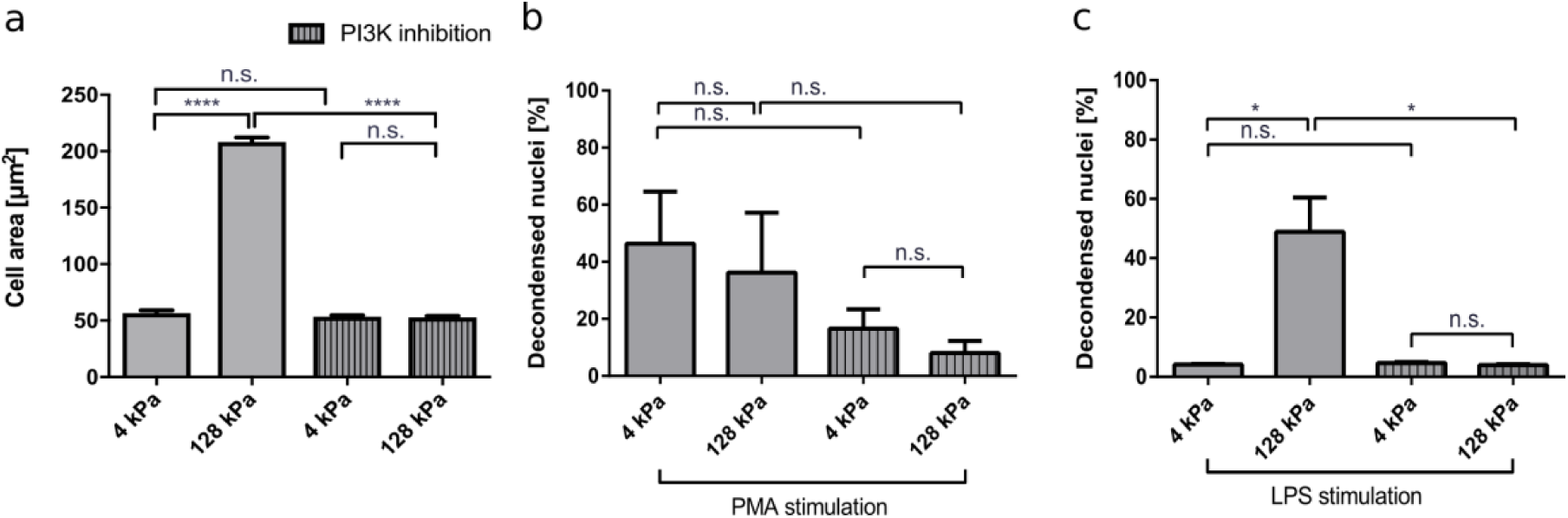
PI3K inhibition impairs neutrophil adhesion and affects NETosis in a stimulant-dependent manner. Human neutrophils were seeded on fibrinogen-coated PAA gels (4 kPa and 128 kPa) and glass (serving as a control) and the spreading area was quantified with and without PI3K inhibition after 30 minutes (a). PI3K inhibition leveled out the elasticity-dependent differences in cell contact area. Then NETosis was induced by PMA (5 nM) (b) or LPS (75 μg/mL) (c) and quantified after 3h. LPS-induced NETosis is completely abolished. In contrast, PMA-induced NETosis is only partially reduced by PI3K inhibition. n > 500 cells per condition. N = 3 donors. Mean ± SEM. Statistics: one-way ANOVA (Bonferroni’s multiple comparisons test. * p < 0.05; ** p < 0.01; **** p < 0.0001; ns: not significant)

To further prove the importance of adhesion for NETosis, we coated glass surfaces with poly-L-lysine (PLL) and poly-L-lysine-grafted-polyethylene glycol (PLL-g-PEG). PEG functionalization/passivation is well established to prevent unspecific adsorption of proteins and adhesion of cells^28^. This environment therefore does not provide any adhesive cues and can be used to test how adhesion affects NETosis. First, cells were seeded on glass, PLL-coated and PLL-g-PEG-coated surfaces and imaged by using reflection interference contrast microscopy (RICM). Dark regions in RICM images indicate close proximity between the cell and the substrate (i.e. adhesion), while bright regions indicate non-adhesive sitting of the cell on the substrate. Cells adhered to both glass and PLL-coated surfaces (**figure 6a, b**). However, they did not adhere to PLL-g-PEG coated surfaces as expected (**figure 6c)**. Cells were stimulated on these coatings and NETosis was found to be independent from adhesion under PMA stimulation as indicated previously^3^ **(Figure 6d).** On the other hand, LPS-induced NETosis was completely inhibited on PLL-g-PEG.

**Figure 6.**
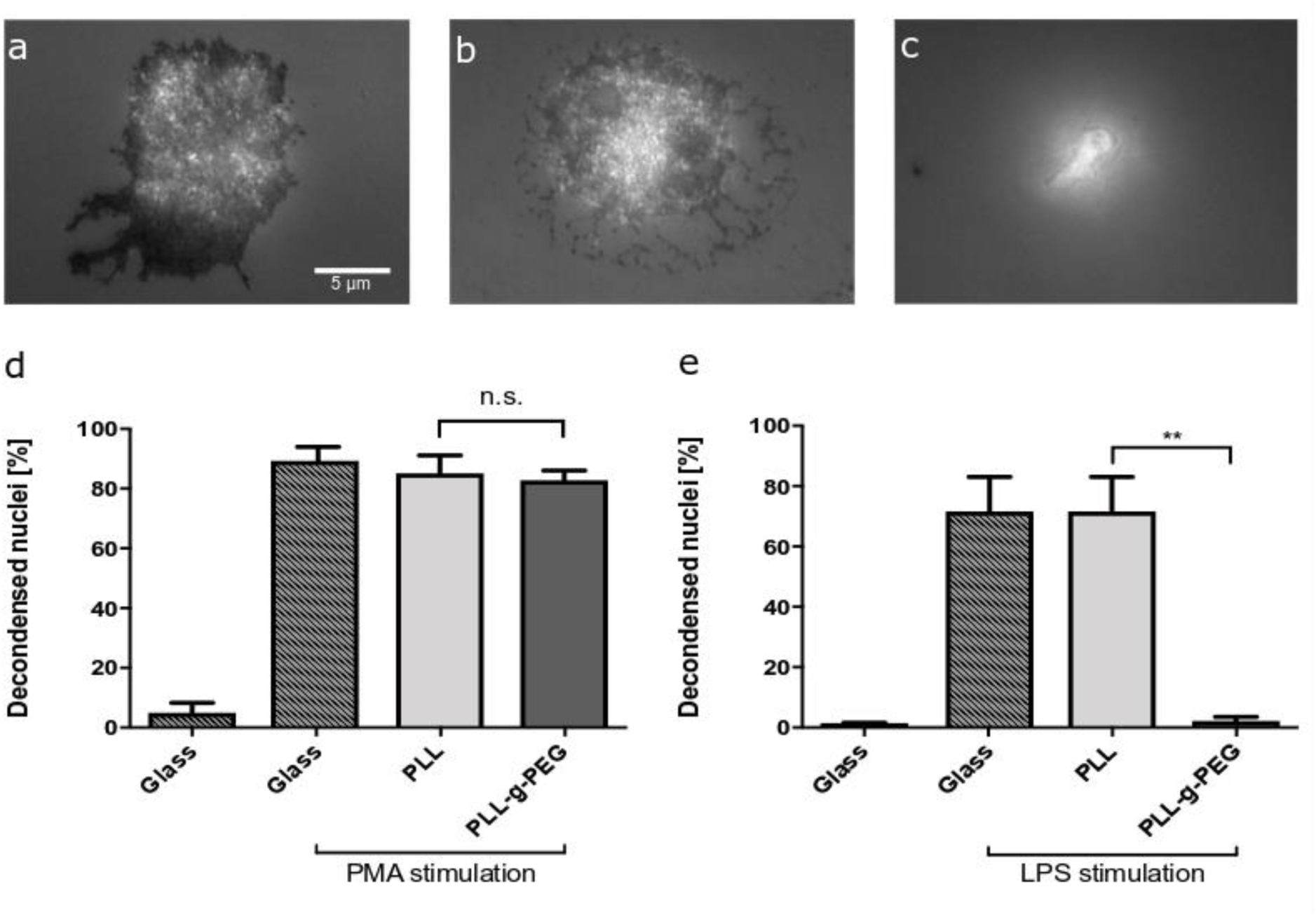
LPS-induced NETosis but not PMA-induced NETosis requires adhesion. RICM images of fixed cells cultured on glass (a), PLL (b) and PLL-g-PEG (c) coated glass surfaces. Compared to glass and PLL surfaces, neutrophils cultured on PLL-g-PEG show no adhesion (c). NETosis on these different substrates was quantified for PMA (d) and LPS stimulation (e). PMA-induced NETosis also took place on the passivated non-adhesive PLL-g-PEG surfaces, while LPS-induced NETosis did not occur. N = 3 donors. Statistics: One-way ANOVA (Bonferroni’s multiple comparisons test ** p < 0.01; ns: not significant). Mean ± SEM.

## Discussion

Neutrophils are the most abundant leukocytes in humans. Being the first responders to inflammation, neutrophils infiltrate different kinds of tissue and subsequently face different mechanical and chemical environments. Substrate elasticity has been shown to influence important cellular functions such as adhesion, differentiation and migration of many different types of cells including neutrophils^10–12,35,41^. Therefore, it is important to understand how NETosis is affected by adhesion in general and more specifically by substrate elasticity. Our results clearly indicate that LPS-mediated NETosis depends on the cells’ adhesion and correlates with the adhesion/contact area, which itself correlates with stiffness.

NETosis is initiated and transmitted *via* diverse pathways, highly depending on the respective stimulus. PI3K is part of the intermediate signaling in PMA-induced and platelet-induced NETosis^37^. PI3K also plays an important role in neutrophil mechanosensing and has been shown to be required for neutrophils to sense substrates of higher stiffness^16^. In order to investigate the role of mechanosensing on neutrophil spreading as well as on NETosis, we tested a highly selective PI3K inhibitor (BAY 80-6946, copanlisib). Indeed, PI3K inhibition impaired spreading of neutrophils and subsequently LPS-induced NETosis. It also inhibited PMA-induced NETosis to a certain degree, which is understandable as PI3-kinase plays a role in PMA-mediated activation of neutrophils. However, it did not abrogate NET formation completely as it did after LPS-stimulation.

Another factor affecting cell adhesion is the available surface concentration of integrin ligands^23,38^. By using nanotechnology approaches it is possible to control the exact distance and overall density of integrin ligands such as RGD and the Mac-1 ligand GPB1α or even link them to advanced near infrared fluorescent nanomaterials^28,42,43^. For neutrophils, adhesion maturation and cellular functions such as spreading and migration depend on ligand density^28^. Here, we showed that on collagen, LPS-induced NETosis not only depends on substrate elasticity but also on the amount of available surface cues. On a substrate that provided a very high density of surface cues (collagen I) cells did not adhere very well and consequently NETosis did not take place. For PMA, NETosis was once again independent of surface cues, in this case density of surface receptors. Indeed, PMA-induced NETosis does not appear to require adhesion at all (**figure 6**). These differences between the stimulants can be explained, at least to some extent, by the different receptor and signaling pathways involved in PMA- and LPS-stimulated NETosis. PMA acts intracellularly and directly activates protein kinase C (PKC). It also triggers subsequent production of reactive oxygen species (ROS), which then interact with MEK, ERK, PI3K, mTOR, MPO and NE^2,44,45^. LPS also activates PKC in neutrophils, primarily through binding to toll-like receptor 4 (TLR4)^46^. Our results show that there is an active connection between adhesion and stiffness signaling and LPS-triggered NETosis, putatively by a connection between integrin and TLR4 signaling, which remains to be explored in depth in the context of NETosis. This hypothesis is also corroborated by the complete abrogation of LPS-induced NETosis following phosphatidylinositol 3-kinase (PI3K) blocking. PMA-induced NETosis is also decreased after PI3K inhibition, thus indicating a role of PI3K in PMA-signaling. However NETosis still took place, albeit at a lower level. It is important to stress that the medium conditions of NETosis assays affect absolute NETosis rates and might shift certain thresholds as previously shown^47^.

Our results might bear considerable medical and pharmacological implications as neutrophils and other cells of the immune system are continuously confronted with surfaces of different stiffness/elasticity. One may speculate that reducing neutrophil responses on and in tissues with low substrate elasticity such as the brain or most importantly the blood itself may serve to keep aberrant immune responses in check. On the other hand, alterations of tissue stiffness as seen in arteriosclerotic vessels, tumor tissue or organ fibrosis might lead to an increase of inflammatory NETosis-related processes^48,49^. Indeed, NET-induced inflammation has been implicated in fibrotic organ changes and is likely to trigger tissue remodeling and lead to even more tissue stiffness, causing a pro-inflammatory, self-sustaining vicious circle^50^. Our results may also be of importance to the explanation of how NETosis can be triggered by implant materials^51^, which tend to have a higher stiffness than “natural” tissues within the body. An improved understanding of the environmental factors that affect NETosis could therefore lead to novel therapeutic approaches for diseases that coincide with alterations of tissue elasticity such as arteriosclerosis, lung or liver fibrosis or cancer. They will also be of great importance for the design of future implant and exogenous materials, which are designed to remain in the body for a certain time, such as catheters.

## Conclusion

In summary, we show how NETosis rates are affected by different levels of adhesion. Neutrophil adhesion increases with substrate stiffness, which leads to higher LPS-induced NETosis rates on stiff substrates. In contrast, PMA induced NETosis does not require any adhesion at all.

## Acknowledgements

This project was supported by the state of Lower Saxony (life@nano) and the German Research Foundation (DFG grant KR 4242/4-1 and ER 723/2-1). Part of this work was supported by the Cluster of Excellence and DFG Research Center Nanoscale Microscopy and Molecular Physiology of the Brain (CNMPB). F.R. acknowledges funding through the DFG within the SFB 937 project A13. We thank Andreas Janshoff and Claudia Steinem for fruitful discussions and support. We are grateful for fruitful discussions about active matter with members of the collaborative research center SFB 937 funded by the DFG.

## Author contributions

S.K and L.E. designed the study. A.L.G., G.G., D.M., E.N., J.G. and L.E. performed NETosis experiments. L.E., A.L.G., M.P.S. and S.K. analyzed data. G.K. and F.R. prepared and characterized gels. L.E., A.L.G., G.G. and S.K. wrote the manuscript with inputs from all authors.

## Data availability

The datasets generated during and/or analyzed during the current study are available from the corresponding author on reasonable request.

## Additional Information

**Supplementary information** accompanies this paper and is available at…

