## Supplementary Information for "Effect of Adhesion and Substrate Elasticity on Neutrophil Extracellular Trap Formation"

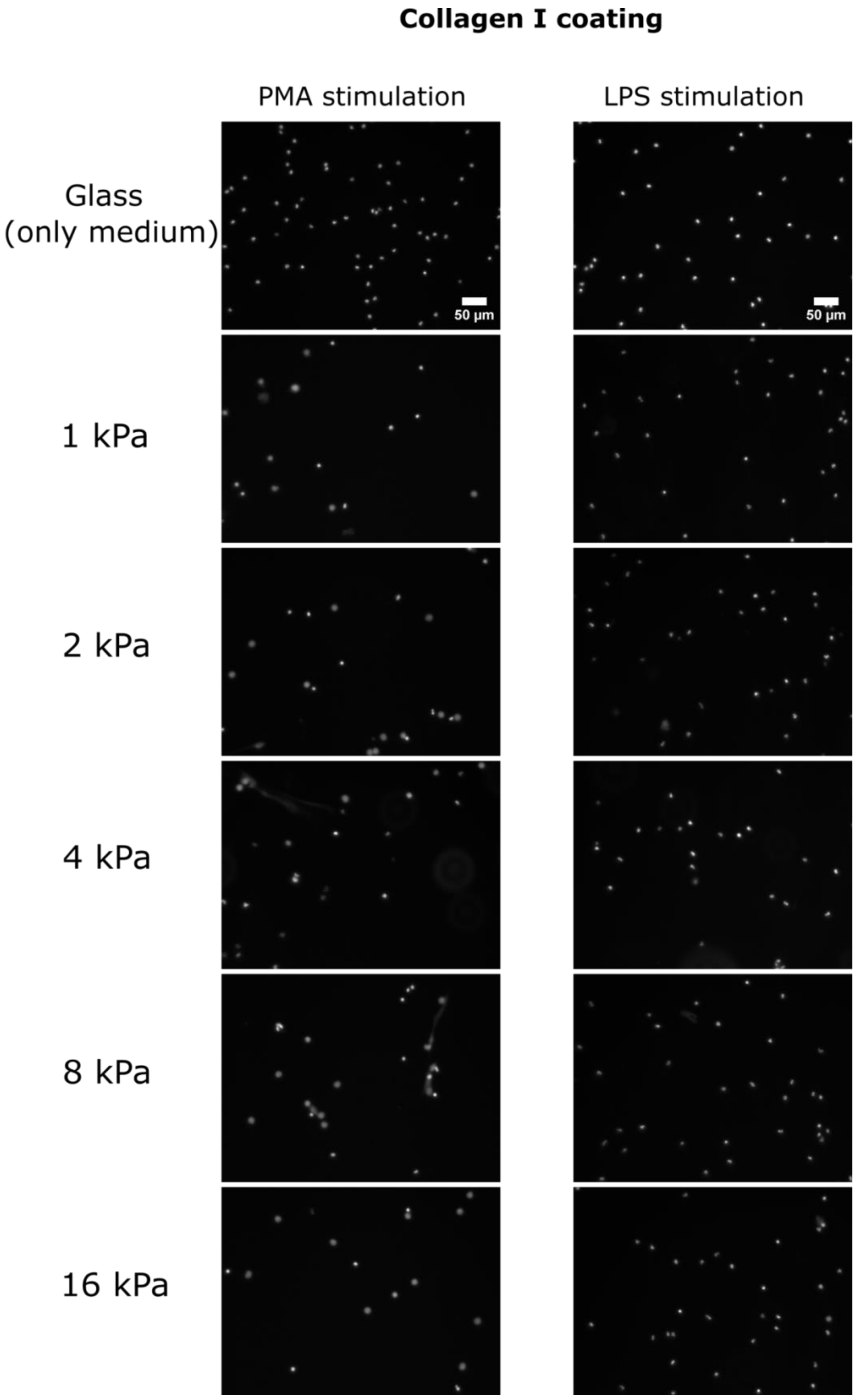

Collagen I coating
PMA stimulation
LPS stimulation
Glass
(only medium)
1 kPa
2 kPa
4 kPa
8 kPa
16 kPa
50 µm
50 µm

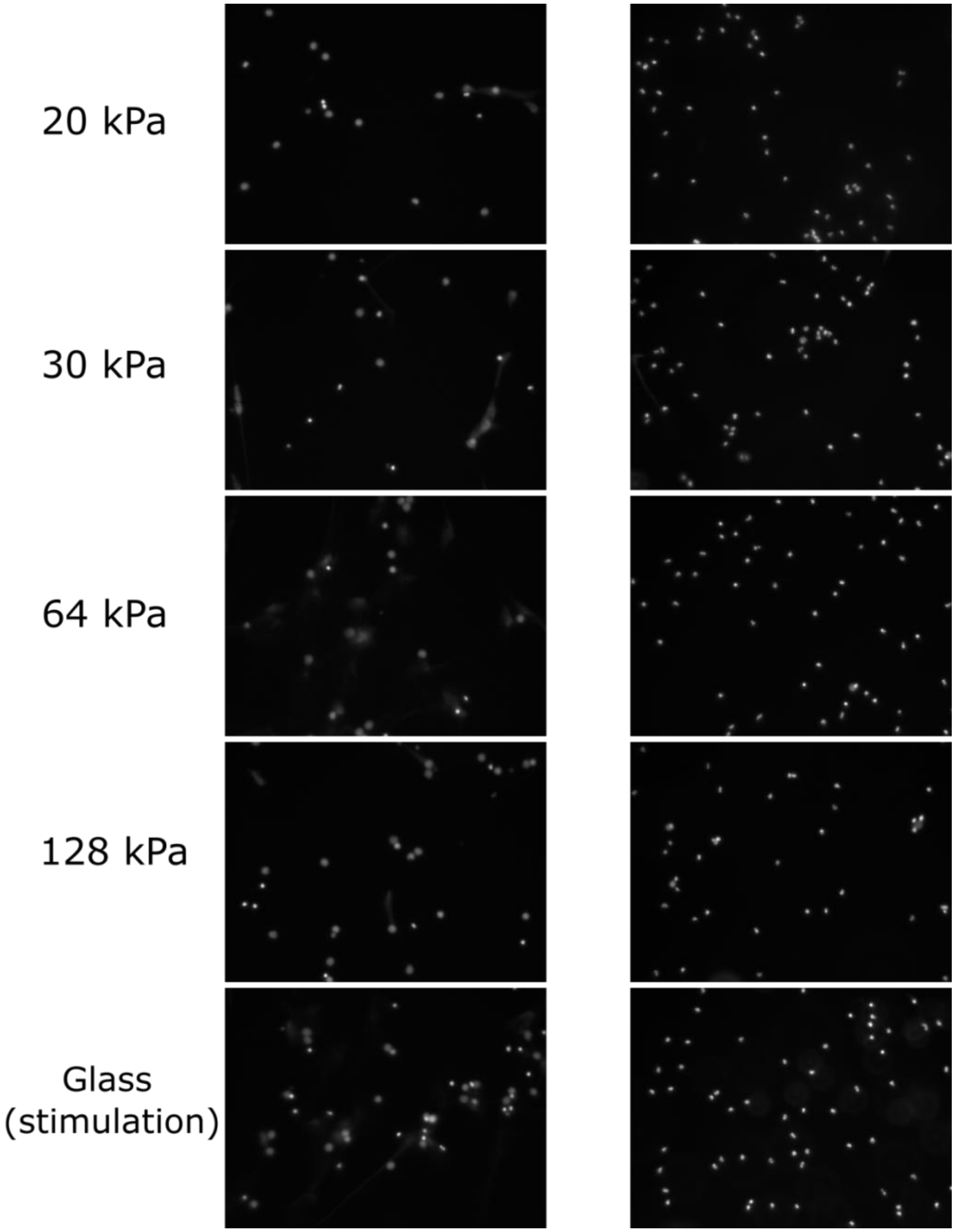


**Supplementary figure S1.** ***Fluorescence images of stimulated neutrophils on PAA gels coated with collagen I.*** *Freshly isolated neutrophils were seeded on polyacrylamide substrates and glass surfaces coated with collagen I and stimulated either with PMA or LPS for 3h. After fixation with 2% PFA, cells were stained with Hoechst 33342/ DNA and representative images were taken from each well to quantify NETosis rates.*

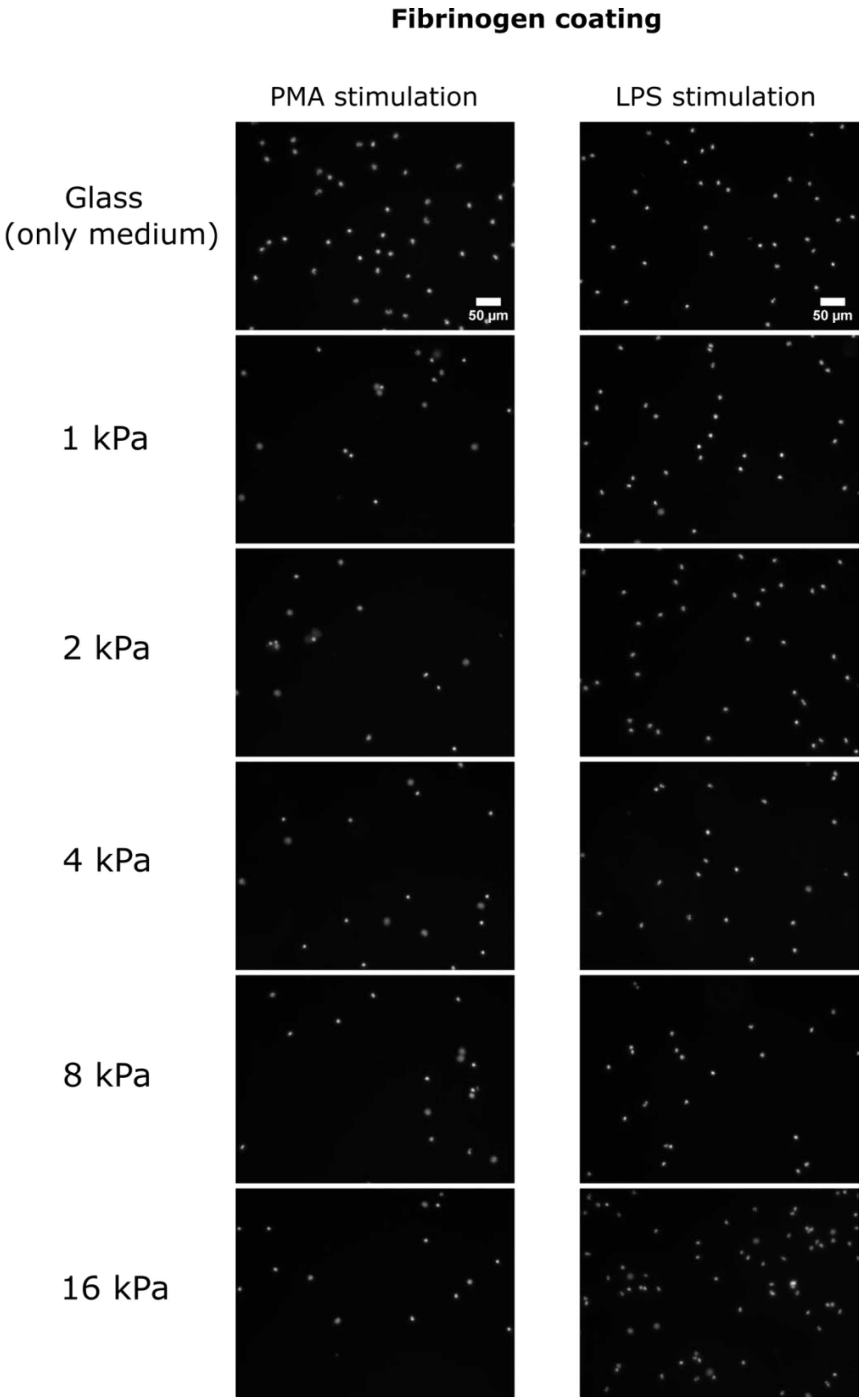
Fibrinogen coating
PMA stimulation
LPS stimulation
Glass
(only medium)
1 kPa
2 kPa
4 kPa
8 kPa
16 kPa
50 µm
50 µm

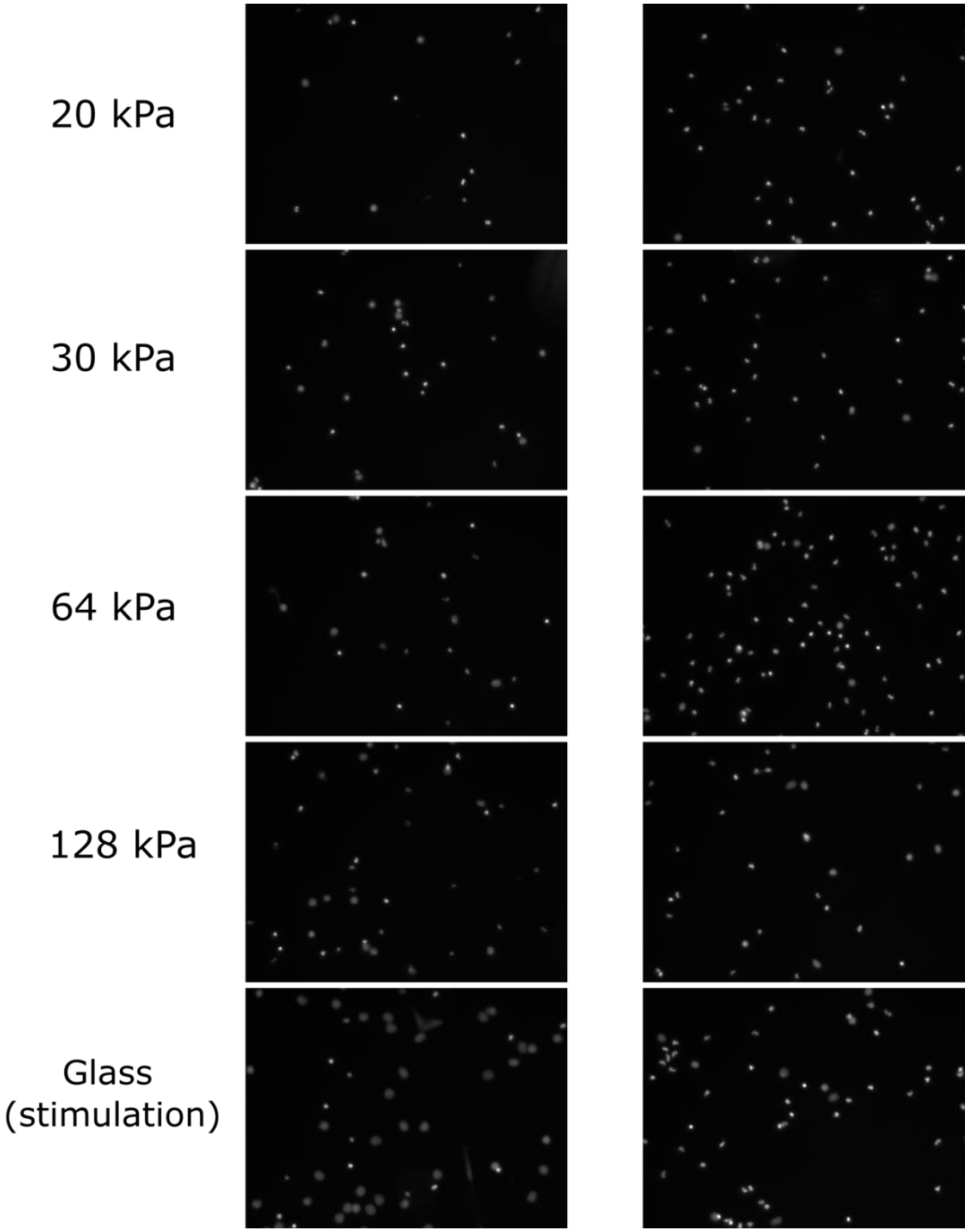


**Supplementary Figure S2** ***Fluorescence images of stimulated neutrophils on PAA gels coated with fibrinogen*****.** *Freshly isolated neutrophils were seeded on polyacrylamide substrates and glass surfaces coated with fibrinogen and stimulated either with PMA or LPS for 3h. After fixation with 2% PFA, cells were stained with Hoechst 33342/ DNA and representative images were taken from each well to quantify NETosis rates.*

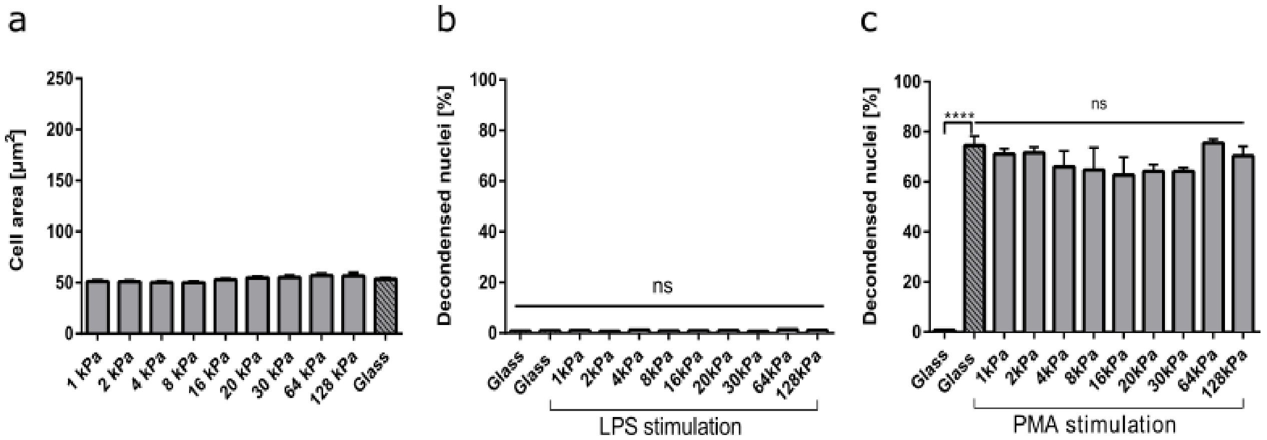


**Supplementary figure S3: NETosis on PAA gels functionalized with higher collagen concentrations.** PAA gels of different elasticity were functionalized with a tenfold higher collagen I concentration (0.2 mg/ml). On those gels neutrophils did not properly adhere and spread (a). Consequently, LPS-induced NETosis did not occur (b). In contrast, PMA-induced NETosis did occur to the same extent on all surfaces.

**Table T1: Composition of the PAA gels and Young's modulus**

| Young's modulus *E* [kPa] | %(v/v) acrylamide in PBS | % (v/v) bis-acrylamide in PBS |
|---|---|---|
| 1 | 3 | 0.20 |
| 2 | 3.5 | 0.20 |
| 4 | 3.8 | 0.20 |
| 8 | 6.8 | 0.10 |
| 16 | 6.8 | 0.20 |
| 20 | 8 | 0.14 |
| 32 | 8.6 | 0.30 |
| 64 | 13.2 | 0.30 |
| 128 | 23.6 | 0.30 |
